# A Cluster of Acidic Residues in the Cytoplasmic Domain of SARS-CoV-2 Spike is Required for Virion-Incorporation and Infectivity

**DOI:** 10.1101/2025.05.04.652125

**Authors:** Charlotte A. Stoneham, Rajendra Singh, Amalia De Leon, Petra Tafelmeyer, Francisco Acosta, Angus Fuori, Michael Anderson, Peter W. Ramirez, Hannah S. Schwartzer-Sperber, Satish Pillai, Mary K. Lewinski, John Guatelli

## Abstract

Like all coronaviruses, the infectivity of SARS-CoV-2 virus particles (virions) requires incorporation of the Spike glycoprotein. Yet, the mechanisms that support the virion-incorporation of Spike are incompletely defined. We noted an unusual feature of human sarbecovirus Spike proteins: their cytoplasmic domains (CDs) contain a stretch of acidic amino acids (DEDDSE). This sequence resembles a cluster of acidic residues, or acidic cluster (AC) motif, found in the cytoplasmic domain of the cellular endoprotease Furin. In Furin, the acidic cluster acts as a protein sorting signal, supporting its intracellular localization at the *trans*-Golgi network (TGN). We tested the contribution of the acidic cluster motif in the Spike CD to protein interactions and to the infectivity of SARS-CoV-2. We used virus-like particles (VLPs) as a model for viral “infection” (transduction). The SARS-CoV2 VLPs were produced by co-expressing Spike (S), Membrane (M), Envelope (E) and Nucleocapsid (N) proteins and deliver an RNA encoding luciferase to target cells expressing the ACE2 receptor. Remarkably, when all five acidic residues of the DEDDSE sequence were replaced with alanines, the VLPs were rendered non-infectious. The N-terminal DE residues provided most of the activity of the acidic cluster. These virologi-cally-impaired Spike mutants were able to reach the cell surface and induce the formation of syncytia, indicating that they are fusogenic and capable of anterograde traffic through the biosynthetic pathway to the plasma membrane. Despite this, they failed to efficiently incorporate into virions. We observed acidic cluster motif-dependent interactions of the Spike CD with several cellular proteins that could potentially support its role in virion-incorporation, including the ERM proteins Ezrin, Radixin, and Moesin; the retromer subunit Vps35, and the medium subunits of the clathrin adaptor complexes AP1 and AP2. While the key cofactor and mechanism of action remains to be defined, this region of acidic residues in the Spike CD appears to be a novel determinant of SARS-CoV-2 infectivity.

## Introduction

The inclusion of the Spike (S) glycoprotein in virus particles (virions) is essential for the infectivity of coronaviruses, but the mechanisms by which S accumulates at sites of viral assembly and incorporates into virions are incompletely understood [reviewed in (*1*)]. These mechanisms likely rely on sequences in S that direct its association with specific membrane domains, on membrane protein trafficking signals in S, and on sequences that specify the recruitment of S into virions via interaction with other proteins, either cellular or viral.

The cytoplasmic domain (CD) of S is the major driver of virion-incorporation; motifs within the CD are responsible for intracellular transport and incorporation of S (Figure 1) (*2, 3*). The membrane proximal, N-terminal half of the SARS-CoV-2 CD is cysteine-rich. Some of these cysteines are palmitoylated, and this modification is needed for efficient virion-incorporation of S and viral infectivity, probably because it directs S into membrane domains where virions assemble (*4, 5*). A region of acidic residues is located centrally within the S CD. This region reportedly supports interactions with several cellular proteins including members of the ERM family (Ezrin, Radixin, and Moesin), the COP-II vesicle coat proteins (Sec23, Sec24), and the retromer complex (SNX27), among others (*6*). These proteins could support the movement of S within the cell in a manner that facilitates virion-incorporation: the ERM proteins via linkage to actin, COP-II vesicles via ER-export, and retromer proteins via retrieval of S to the *trans*-Golgi network (TGN). At the C-terminus of the CD, a lysine-based ER-retention signal is present (KLHYT) (*7*). This sequence binds COP-I vesicle coats, which mediate retrograde traffic from the Golgi to the ER (*8*), but it does so inefficiently, allowing S to escape the ER and reach the ER-Golgi intermediate compartment (ERGIC) (*6*), where virions assemble (*9*).

**Figure 1.**
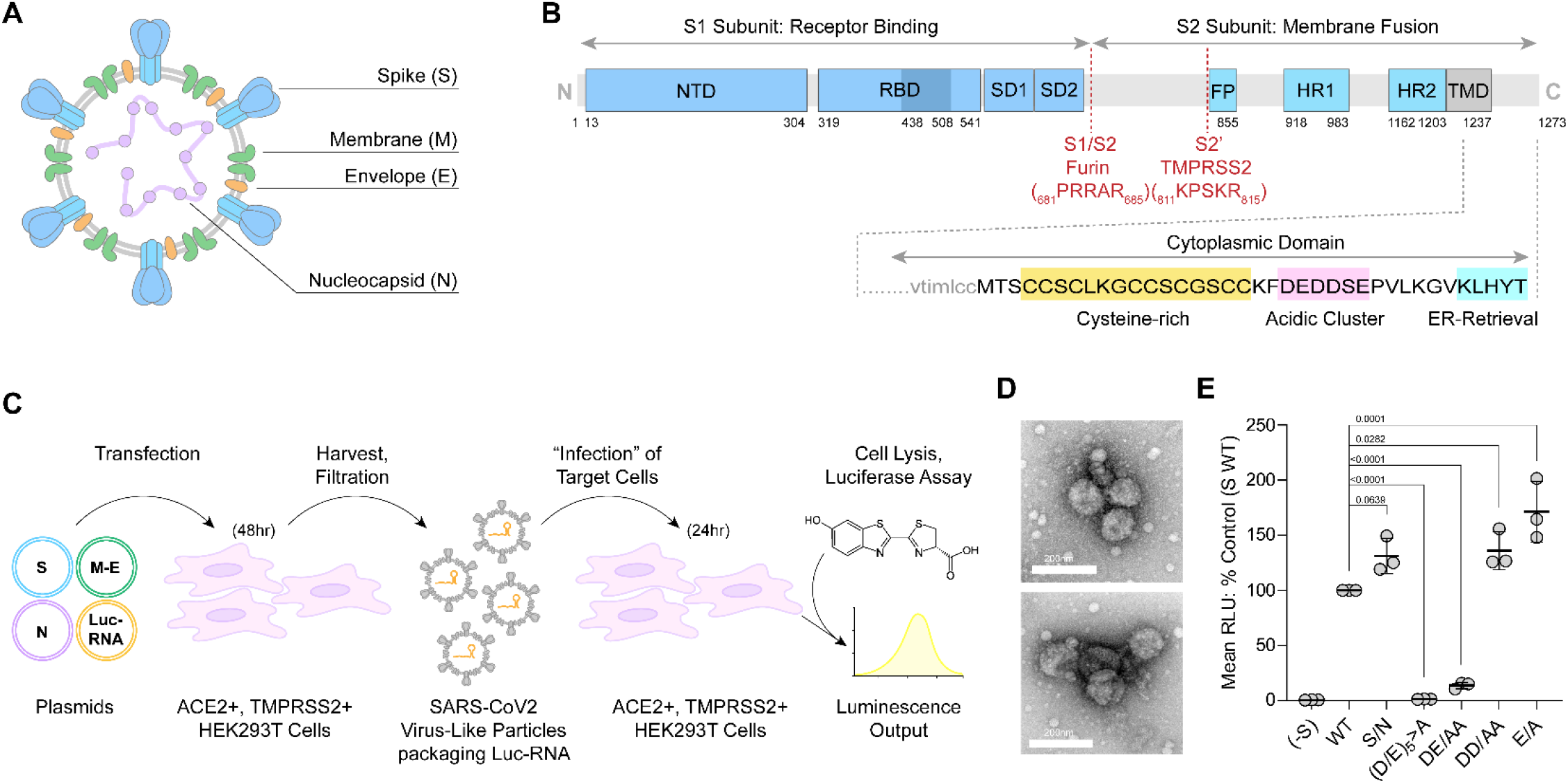
Infectivity (transducing activity) of authentic SARS-CoV-2 virus-like particles (VLPs) requires the acidic cluster in the CD of S. **A.)** Organization of SARS-CoV2 virion major structural proteins and **B.)** the Sequence of the cyto-plasmic domain of S. The cysteine-rich domain, the acidic cluster motif, and the ER-retention signal are highlighted. **C.)** Schematic of the authentic SARS-CoV-2 VLP luciferase transduction system (*20*). We modified the published system by using HEK293T cells that express ACE2 and TMPRSS2 as both the VLP producer cells and as the targets of infection (transduction) by the VLPs. S: Spike. M-E: Membrane-IRES-Envelope. N: Nucleocapsid. Luc-RNA: RNA encoding luciferase and containing a packaging determinant from the N coding sequence. **D.)** Electron micrographs of negatively-stained preparations of SARS-CoV2 VLPs; particles of approximately 100nm size were identified in crude preparations, which appear decorated with Spike glycoprotein **E.)** Relative infectivity of SARS-CoV-2 VLPs measured as expression of transduced luciferase activity (relative light units; RLU). Data are from three independent experiments and are normalized to wild type (“WT”) S as 100%. “(D/E)_5_>A” denotes substitution of the 5 acidic residues for alanines, “DE/AA” is a mutant of the N-terminal acidic residues, “DD/AA” is a mutant of the central residues, and “E/A” is a mutant of the C-terminal E of the sequence. *P*-values are derived from one-way ANOVA analysis with Dunnett multiple comparisons correction; comparing mutants to WT control.

We noted that the cluster of acidic residues in the S CD – DEDDSE-resembles the acidic cluster motifs found in the CDs of other cellular and viral proteins, including the endoprotease Furin, which processes viral glycoproteins including SARS-CoV-2 S (*10-13*). Phosphorylation of serine in the Furin acidic sequence (SDSEEDE) creates an extended negative-charge region that binds the positively charged medium subunits of clathrin adaptor protein (AP) complexes (*12, 14*). These interactions direct Furin to the *trans*-Golgi region (*10*). Simlarly, the acidic cluster in the S CD could be an AP-binding signal that concentrates S at sites of viral assembly. Comparative virology supports this: the CDs of S proteins from diverse coronaviruses contain a Yxxφ motif (where φ is bulky hydrophobic residue) (*1*), and Yxxφ sequences bind the medium subunits of clathrin AP complexes (*15*). SARS-1 and SARS-CoV-2 lack a Yxxφ sequence. Plausibly, the DEDDSE sequence is an alternative clathrin AP-binding sequence utilized by these sarbecoviruses.

In addition to cellular proteins, virus-encoded proteins are candidates for mediating the recruitment of S into virions [reviewed in (*1*)]. For example, interactions between S and M are well documented in several coronaviruses (*16-18*) . Relevant to the potential for an S-M interaction, cryo-EM structures of the M protein of SARS-CoV-2 reveal basic patches on the cytoplasmic surface of SARS-CoV-2 M protein (*19*); these are plausible partners of the acidic cluster in the S CD.

Here we show that the acidic cluster in the S CD is essential for the infectivity of virus-like particles (VLPs) made from authentic SARS-CoV-2 structural proteins, and it is essential for the incorporation of S into those particles. We confirm several cellular proteins as acidic cluster-dependent interactors of the S CD and extend that list to include the medium subunits of the clathrin adaptor complexes AP1 and AP2.

## Results

### An authentic SARS-CoV-2 VLP assay reveals the essential nature of the acidic cluster in the CD of S for infectivity

SARS-CoV-2 virus-like particle (VLP) systems are facile, reductive platforms with which to probe the determinants of viral assembly and infectivity without the need to manipulate infectious virus. We used a previously reported SARS-CoV-2 VLP system in which the structural proteins Spike, Membrane, Envelope, and Nucleocapsid are co-expressed in virus producer cells, along with a packagable mRNA reporter encoding luciferase (*20*). Figure 1 explains this system and shows that residues of the acidic cluster in the CD of S are required for viral infectivity. As expected, virtually no infectivity (transduced luciferase activity) is detected in the absence of S (“-S’). Substitution of the acidic resides of the acidic cluster with alanines (_1257_DEDDSE_1263_/AAAASA, “(D/E)_5_>A”) eliminated infectivity. Substitution of the N-terminal DE of the DEDDSE (“DE/AA”) sequence reduced infectivity by 80%. Other substitutions in the sequence, including the S/N mutation, did not impair infectivity.

### The acidic residues of the DEDDSE sequence are dispensable for the formation of syncytia

We checked Spike function in the above experiments independently of VLP-infectivity by observing the producer (transfected) cells for syncytia (Figure 2). Large syncytial cells were not observed in the absence of S, but they were observed when either wild type S, S-(D/E)_5_>A, or S-DE/AA was expressed. The DD/AA, S/N, and C-terminal E/A mutants also formed syncytia indistinguishably from the WT (data not shown for the S/N mutant).

**Figure 2.**
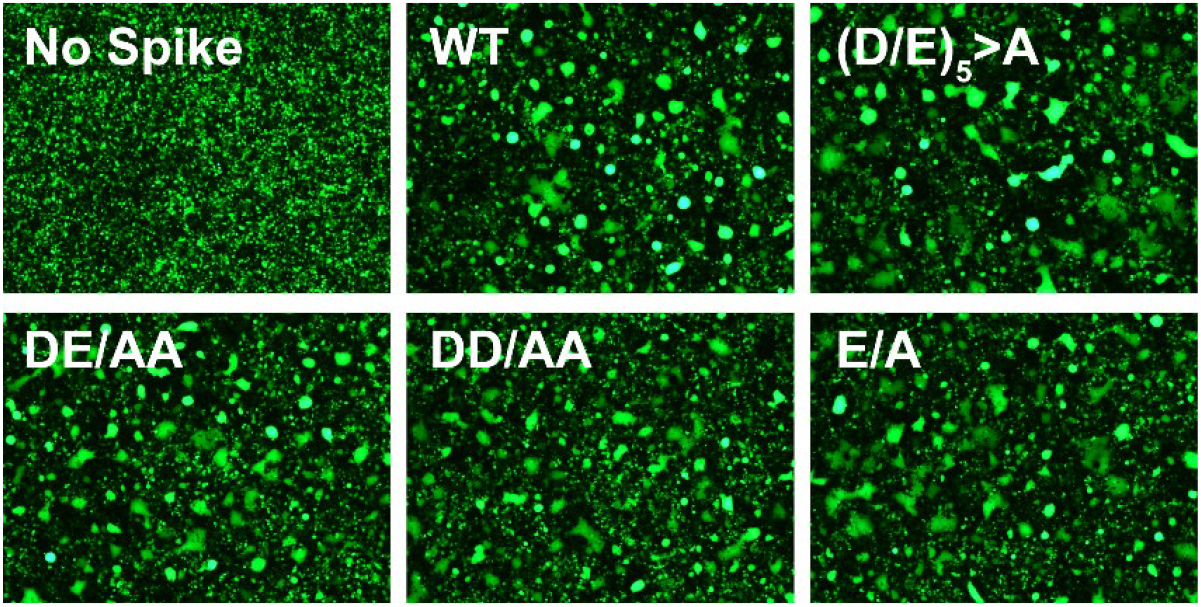
Mutation of the acidic residues does not impair formation of syncytia. The formation of syncytia was monitored by including a plasmid expressing GFP in the transfection of the cells that produced the VLPs, which express both ACE2 and TMPRSS2. Images of live cells were obtained 48 hours after transfection with the S, N, and M-IRES-E plasmids, just before the VLPs were harvested. The same ability to induce syncytia was observed in parallel experiments in which S was expressed in isolation, i.e., without co-expression of M, N, and E (not shown).

The key conclusions from the microscopic data are that 1) lack of fusogenicity does not explain the inability of alanine-substituted acidic residue mutants (D/E)_5_>A and DE/AA S to support infectivity and 2) the (D/E)_5_>A and DE/AA S mutants reach the cell surface (also confirmed by flow cytometry; data not shown). To have reached the cell surface, they presumably traversed the ERGIC, where virion-assembly occurs.

### Acidic residues of the DEDDSE acidic cluster are required for the efficient incorporation of S into VLPs

Figure 3 shows that the acidic cluster directs the incorporation of S into VLPs. Abundant S protein was detected in pelleted VLPs made with wild type S. In contrast, VLPs made with the (D/E)_5_>A or DE/AA mutants incorporated very little S. The lack of virion-incorporation was not due to a failure of expression (see the “Cell Lysate” panel of Figure 3).

**Figure 3.**
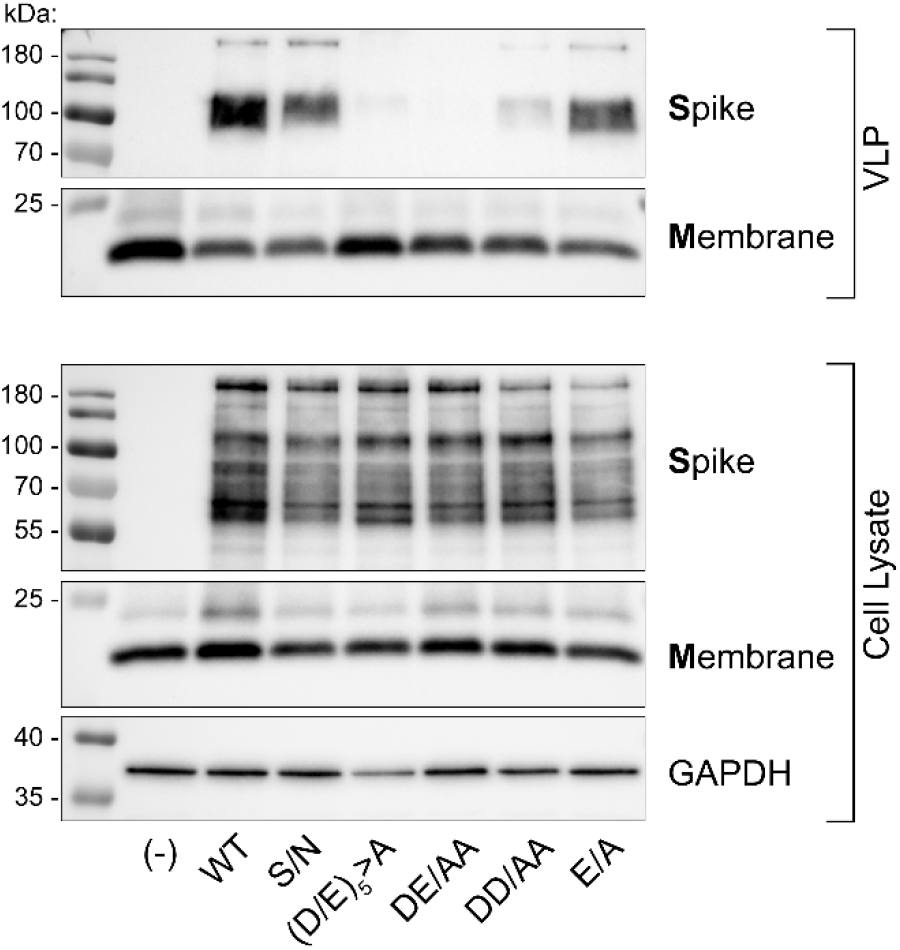
Virion incorporation of S requires the acidic cluster and depends largely on the N-terminal DE residues. VLPs from one of the experiments shown in Figure 2 were analyzed by western blot, as were the cells that produced them. VLPs were partially purified from the cultures by pelleting out cellular debris followed by filtration through 0.2 micron membranes, and then pelleting at 23,500 x *g* for 2 hours. *Top:* Pelleted material from the VLP preparations (“VLP”) were analyzed by immunoblots probed with antibody to Spike as well as Membrane protein, a key structural component of virions. *Bottom:* aliquots of the VLP producer cells (“Cell Lysate”) were similarly probed for S, M, and the cellular protein GAPDH. VLP-associated S included a minor amount of high molecular weight unprocessed S but was prominently a doublet around 100 kDa, which is likely S processed by Furin and TMPRSS2(*21*, *22*). VLPs made with the S DEDDSE/AAAASA ((D/E)_5_>A) and DE/AA mutants were nearly devoid of S signal, whereas VLPs made with S DD/AA have diminished signal. VLP producer cells show similar amounts of WT and mutant S proteins without obvious differences in band-patterns.

The DD/AA mutant S incorporated into VLPs more efficiently than the (D/E)_5_>A and DE/AA mutants but substantially less efficiently than the WT. Yet, its infectivity was unimpaired (compare Figures 1 and 3). While these data on the DD/AA mutant can be interpreted to weigh against the hypothesis that impaired virion-incorporation explains the infectivity phenotypes of these mutants, they can also be interpreted as indicating a non-linear relationship between the incorporation of S into the VLPs and their infectivity, such that the infectivity measured requires a threshold of incorporated S reached by the DD/AA mutant (and the S/N and E/A mutants).

### Acidic cluster-mediated protein interactions of the S CD

We evaluated potential cellular protein-partners of the acidic cluster based on our unpublished two-hybrid screen, published data(*6*), and the similarity between the S acidic cluster and the phospho-serine acidic clusters found in cellular proteins such as Furin(*10-12*).

### Interactions of the S CD detected by yeast two-hybrid assays: ERM proteins

Figure 4A shows the results of yeast two-hybrid assays using the S CD as “bait” and specific cellular proteins as “prey.” The S CD interacted robustly with Ezrin, HTRA2, Radixin, Moesin, LAMA1, and RBBP5. Very weak potential interactions with α-COP, a subunit of the COP-I vesicle coat and with vps35, a subunit of the retromer complex, were also detected. Of these, the interactions with Ezrin, Radixin, Moesin, RBBP5, and possibly α-COP and Vps35, but not HTRA2 and LAMA1, required the acidic residues of the acidic cluster.

**Figure 4.**
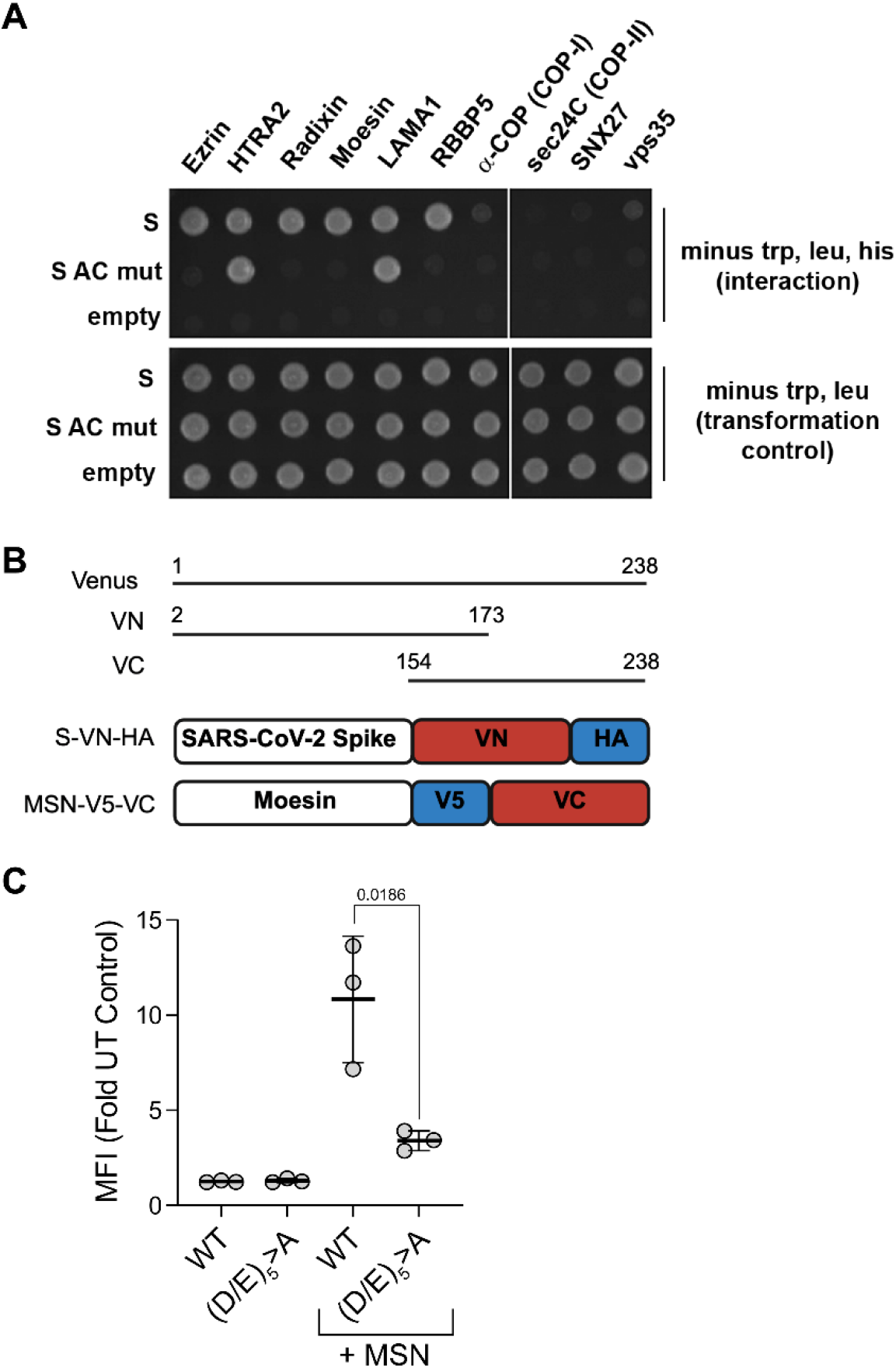
Yeast two-hybrid and Bi-molecular Fluorescence Complementation (BiFC) protein-protein interaction assays show acidic cluster-dependent interactions of the S CD with various cellular proteins. **A.)** *Top panel*: yeast growth in the absence of tryptophan, leucine, and histidine indicates interaction between the “prey” proteins along the top and the “bait” proteins at the left. “S” indicates the wild type S CD; “S AC mut” indicates the (D/E)_5_>A S mutant; “empty” indicates empty bait-vector control. *Bottom panel*: yeast growth in the absence of tryptophan and leucine indicates the presence of the bait and prey plasmids. **B.)** BiFC constructs. The Venus protein (residues 1 to 238) is divided into N-terminal (VN: residues 2 to 173) and C-terminal portions (VC: 154 to 238). VN was fused to the C-terminus of SARS-CoV-2 Spike and VC to the C-terminus of Moesin, along with epitope tags (HA or V5) as shown. **C.)** Cells (HEK293T) were transfected with plasmids expressing the indicated proteins, either Spike-WT-VN or Spike-(D/E)_5_>A -VN, with or without co-expression of Moesin (MSN)-VC. Twenty-four hours later, the mean fluorescence intensities (MFI) of the cells were determined by flow cytometry. Results are the mean fold difference in fluorescence relative to untransfected cells. Error bars represent the SD from three biological experiments performed in technical duplicate. *p* value (unpaired t test) between Spike-WT-VN or Spike-(D/E)_5_>A -VN co-expressed with MSN-VC is indicated. Expression of the BiFC fusion proteins was confirmed by western blot (data not shown).

### Interactions of the S CD detected by Bi-molecular Fluorescence Complementation (BiFC): Moesin

We used BiFC to show that the acidic cluster in the S CD supports an interaction between full length Spike and Moesin in living human cells (Figure 4, panels B and C). In BiFC, the proteins of interest are fused to N-or C-terminal fragments of Venus (yellow fluorescent protein (YFP)). Protein-protein interaction reassembles YFP, generating fluorescence (*23*). Spike-VN (Spike fused to the N-terminal fragment of Venus) and Moesin-VC (Moesin fused to the C-terminal fragment of Venus) together yielded a BiFC signal greater than Spike-VN alone, suggesting an interaction. Consistent with our yeast two hybrid data, mutating the acidic residues of the S-CD significantly reduced that signal (Figure 4, panels B and C).

### Interactions between the S CD and the medium (µ) subunits of clathrin adaptor proteins detected by GST-pull-down using recombinant proteins

We used recombinant proteins to look for direct interactions between the S-CD and the µ subunits of AP2, the clathrin adaptor associated with endocytosis of membrane proteins(*24*), and AP1, the clathrin adaptor associated with retrieval of membrane proteins to the TGN(*25*) (Figure 5).

**Figure 5.**
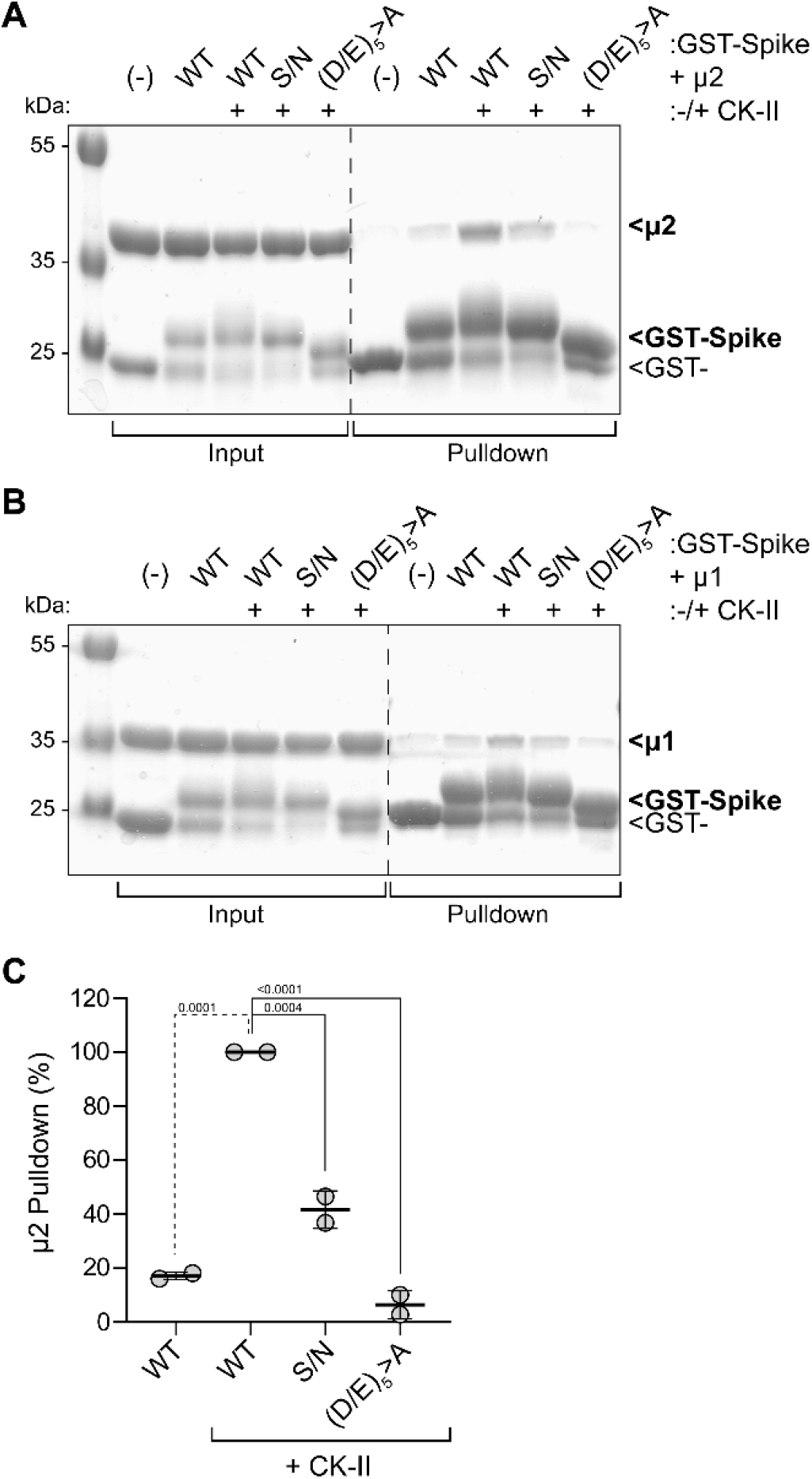
Acidic cluster-dependent interactions of the S CD with the medium (µ) subunits of the clathrin adaptors AP1 and AP2. **A.)** Pulldowns using GST-S-CD and µ2 (the medium subunit of the AP2 adaptor protein complex). **B.)** Same experimental design but using µ1 of the AP1 complex. All proteins were recombinant and produced in *E*.*coli*. The GST-S-CD proteins were expressed either with or without Casein Kinase II (CK-II) to phosphorylate serine residues. Pulldowns were assessed by SDS-PAGE and staining with Coomassie Blue. **C.)** Quantitation of replicate µ2 pull-down experiments by densitometry. *p* values are derived from one-way ANOVA analysis with Dunnett multiple comparisons correction, comparing values to Spike-CD WT (+ CK-II) control.

The S-CD bound both µ1 and µ2. These interactions required co-expression of Casein Kinase II (CK-II), a serine/threonine kinase. The CK-II-dependent µ-interactions required the acidic residues and to a lesser extent the serine of the acidic cluster. These data extend the similarity between the acidic cluster in the S-CD and the phospho-serine acidic clusters found in several cellular and viral membrane proteins, including the prototype sequence SDSEEDE in Furin, which require-serine phosphorylation for µ-binding and to function as sorting motifs(*10, 12*). Notably, the intermediate phenotype of the S/N mutant in µ-binding is consistent with its intermediate phenotype in virion-corporation (Figure 3), although the infectivity of VLPs produced using this mutant is not diminished (Figure 1).

## Discussion

We have made the novel observation that the acidic cluster centrally located within the cytoplasmic domain of SARS-CoV-2 Spike is required for virion-infectivity and virion-incorporation, at least in the setting of an authentic SARS-CoV-2 infectious virus-like particle (VLP) assay. While this sequence has been studied in terms of its interactions with cellular proteins and its impact on the trafficking of Spike (*6*), to our knowledge this is the first report of its key virologic role.

To date, the most compelling cell biologic role for this sequence is its contribution to the interaction of the S-CD with COP-II vesicle coats, which mediate ER-to-TGN transport (*6*). In principle, this fits well with the virologic observations herein: if S cannot exit the ER, it would not be able to reach viral-assembly sites in the ERGIC. However, mutants of the S acidic cluster showed similar banding patterns on immunoblots of denaturing SDS-PAGE gels; this weighs against a failure of ER-exit, which should have caused differences due to incomplete glycosylation. Moreover, we show here that acidic cluster mutants that are virologically defective or severely impaired do reach the plasma membrane, where they are able to cause cell-cell fusion. Absent a novel route of ER exit that leads to the plasma membrane while bypassing the Golgi, these Spike mutants must have exited the ER and moved through viral assembly sites, yet they failed to incorporate efficiently into virions.

What could be the key partner of the acidic cluster that facilitates virion-incorporation and infectivity? Importantly, that partner could be cellular or viral, and more than one protein might be involved. Among cellular proteins reported so far (*6*), we confirm here acidic cluster-dependent interactions of the Spike CD with ERM proteins (Ezrin, Radixin, and Moesin) and potentially a subunit of the retromer complex (Vps35) using yeast two-hybrid and BiFC assays. The yeast two-hybrid assay also identified an interaction with RBBP5, a component of a histone methyltransferase complex (*26*), of unclear significance. We extend the interaction partners of the S CD to include the µ subunits of clathrin adaptor protein complexes (AP2 and AP1). The interactions with µ subunits are direct and require not only the acidic residues but also phosphorylation of the S-CD, at least *in vitro*, by CK-II. The AP complexes and retromer would presumably retrieve S to viral assembly sites (*27*); yet, we have not observed striking changes in the intracellular distributions of native S at steady-state caused by mutation of the acidic cluster (data not shown). On the other hand, the role of ERM proteins in viral assembly is highly speculative: they could provide correct trafficking of S by linkage to actin (*28*), or they could provide a novel scaffolding function that supports virion-incorporation of S. If the acidic cluster mediates a direct interaction with a viral protein, that protein must be either M, N, or E, the only other viral proteins expressed in the VLP assay. One clue to the identity of the acidic cluster’s binding partner is that the interaction is likely electrostatic, involving the negatively charged acidic cluster and a positive charged basic domain on the partner protein. Notably, the protein-partners considered here have such basic domains (Figure 6).

**Figure 6.**
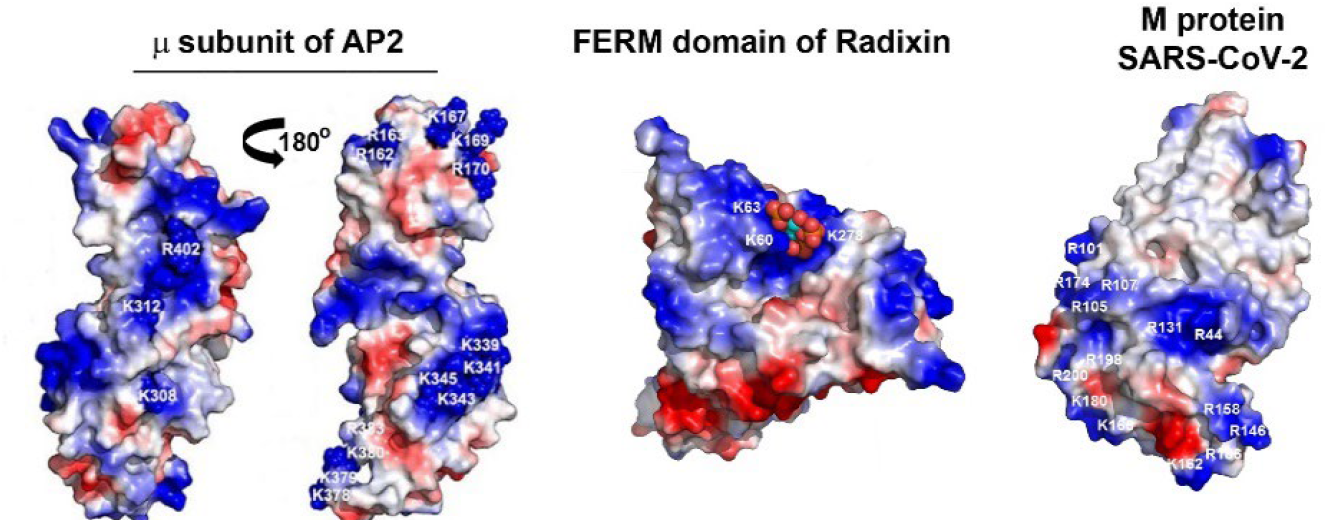
Basic domains on the surfaces of potential protein partners of the S acidic cluster. Surfaces of the C-terminal two-thirds of the µ subunit of AP2 (PDB:1BW8), the FERM domain of Radixin (PDB:1GC6), and the M protein of SARS-CoV-2 (PDB:8CTK). Positively charged regions are colored blue, negatively charged regions are red. The positively charged regions could bind the acidic cluster in the S-CD. Spheres on the FERM domain indicate bound PIP2.

What can comparative virology tell us about the acidic cluster? As noted above, some coronavirus Spike CDs contain Yxxφ sequences that have been suggested to support trafficking or endocytosis(*1*). Yxxφ is the canonical linear interaction motif that binds to the µ subunits of clathrin adaptors (*15*), so we are tempted to speculate that sarbecovirus S, which has no such sequence, uses a phospho-serine acidic cluster for µ-binding instead. Still, why this sequence rather than a Yxxφ sequence? Moreover, does this evolutionary choice mean anything about the species specificity or pathogenesis of sarbecoviruses? Notably, a virologic observation supports a role for the acidic cluster in sending S to the correct intracellular location for virion-assembly: the C-terminal 19 residues of the S CD, which includes the acidic cluster, are typically deleted for the purpose of efficiently pseudotyping retroviral and VSV-vectors with sarbecovirus S (*29, 30*). The site of assembly of these pseudoviruses is the plasma membrane not the ERGIC (*31, 32*). If the acidic cluster drives the accumulation of S at the ERGIC, then it should inhibit the pseudotyping of retroviruses and rhabdoviruses.

The main caveat to our virologic data is that it is limited to a VLP system. While this system uses only authentic SARS-CoV-2 proteins, it has a remarkably strict dependence on the amount of S expression; both too much and too little reduce the yield of infectious VLPs (*20*). Moreover, the stoichiometry of the structural proteins including S expressed in this system is empiric and might not approximate that of viral replication. The introduction of acidic cluster mutations into replication-competent virus and the characterization of such mutants in various cell culture and animal models is needed to validate the findings herein.

In summary, we report here a novel and key role of the S-CD acidic cluster in SARS-CoV-2: it is required for infectivity and virion-incorporation in a model, virus-like particle system. Whether these findings will hold true in the case of the actual virus, and what the key interaction-partners are, remain to be determined.

## Acknowledgements

We thank Marissa Suarez for technical support. This work was supported by grants U19 AI171443 and R16 AI184450 from the NIH, USA, and by the French National Science Foundation (COV2PIM (ANR-20-COVI-0089-01)

## Methods

### Plasmids and mutagenesis

Plasmids expressing wild type SARS-CoV-2 from Wuhan-Hu-1 (pCG1-SARS-2-S) was a gift from Stefan Pöhlman (*21*). Substitutions of acidic residues in the cytoplasmic domain of S were introduced using Quikchange PCR Mutagenesis XL kit (Agilent); _1261_S/N: Fwd 5’-GCAC-GGGCTCATTATCGTCCTCGTCGAACTTGC-3’; Rev 5’-GCAAGTTCGACGAGGACGATAATGAGCCCGTGC-3’, _1257_DEDDSE/AAAASA: Fwd 5’-CCTTCAGCACGGGCGCAGAAGCGGCCGCGGCGAACTTGCAGCAG-3’; Rev 5’-CTGCTGCAAGTTCGCCGCGGCCGCTTCTGCGCCCGTGCTGAAGG-3’, _1257_DE/AA: Fwd 5’-GGCTCAGAATCGTCCGCGGCGAACTTGCAGCAGC-3’, Rev 5’-GCTGCTGCAAGTTCGCCGCGGACGAT-TCTGAGCC-3’, _1259_DD/AA: Fwd 5’-GCACGGGCTCAGAAGCGGCCTCGTCGAACTTG-3’, Rev 5’-CAAGTTCGACGAGGCCGCTTCTGAGCCCGTGC-3’, _1262_E/A: Fwd 5’-TTCAGCACGGGCG-CAGAATCGTCCTCGTCG-3’, Rev 5’-CGACGAGGACGATTCTGCGCCCGTGCTGAA-3’.

Plasmids expressing N-R203M, M-IRES-E, and Luc-PS9 were gifts from Jennifer Doudna (*20*). pCG-GFP was a gift from Jacek Skowronski. Mutagenesis of pCG1-SARS-2-S was done using the Quikchange kit (Agilent Technologies) and custom oligonucleotides (synthesized by Integrated DNA Technologies); mutations were confirmed by Sanger sequencing (Genewiz; Azenta Life Sciences).

Gibson assembly was used to create Spike-VN-HA and MSN-V5-VC. The coding sequences for Spike (CoV2-Spike-EF1a-D614G-N501Y; AddGene #177939), Moesin (Sino Biological; #HG13866), VN-HA and V5-VC (*33*) were PCR-amplified to contain overlapping ends using the following primers: Spike (Forward): 5’-TAGTCCAG-TGTGGTGGAATTATGTTTGTCTTCCTGGTCCT; Spike (Reverse): 5’-AGCTCCTCGCCCTTGCTCACTG-TATAGTGCAGTTTGACGC-3’; VN-HA (Forward): 5’-GCGTCAAACTGCACTATACAGTGAGCAAGGGCGAG-GAGCT-3’; VN-HA (Reverse): 5’-CCCTCTAGACTCGAGCGGCCTCAAGCGTAATCTGGAACATCGT-3’; Moesin (Forward): 5’-TAGTCCAGTGTGGTGGAATTATGCCCAAAACGATCAGTGT-3’; Moesin (Reverse): 5’-TTAGGGATAGGCTTACCCATCATAGACTCAAATTCGTCAATGC-3’; V5-VC (Forward): 5’-TTGACGAATTT-GAGTCTATGATGGGTAAGCCTATCCCTAA-3’ ; V5-VC (Reverse): 5’-CCCTCTAGACTCGAGCGGCCTCAC-TTGTACAGCTCGTCCA. PCR products were ligated into *Kpn* I and *Xho* I linearized pcDNA3.1+ (Invitrogen) using Gibson Assembly Master Mix according to the manufacturer’s recommendations (New England Biolabs), and the assembled DNA was subsequently transformed into DH5-alpha competent *E*.*coli* (New England Biolabs). Plasmid DNA was confirmed by Sanger sequencing. Spike-AAAASA-VN-HA was created using the Q5 Site-Directed Mutagenesis kit according to the manufacturer’s instructions (New England Biolabs) using the following primers: Forward 5’-GCTAGCGCGCCCGTGCTGAAGGGCGTC-3’; Reverse 5’-GGCCG-CAGCGAACTTGCAGCAGCTGCC-3’.

### Cells

HEK293T cells (ATCC: CRL-3216) were grown in Dulbeccco Modified Eagle’s Medium (DMEM) supplemented with 10% Fetal Bovine Serum (FBS; Corning) and 1% Penicillin/Streptomycin (Gibco). HEK293T cells expressing ACE2 and TMPRSS were made as follows. ACE2 was cloned into pLKO5d.SFFV.dCas9-KRAB.P2A.BSD (a gift from Dirk Heckl, Addgene plasmid #90332) by replacing dCAS9-KRAB with new unique enzyme restriction sites (*Spe* I and *Nhe* I) and then inserting the ACE2 gene sequence into the expression construct downstream of the SFFV promoter. TMPRSS2 was cloned into pDUAL CLDN (GFP) (a gift from Joe Grove, Addgene plasmid #86981). GFP was exchanged with a puromycin cassette using *Mlu*l and *Xho*I sites. TMPRSS2 was inserted into the expression construct immediately downstream of the SFFV promoter following the addition of unique enzyme restriction sites (*Srf* I and *Sal* I). Both constructs were confirmed by Sanger sequencing. Lentiviral particles were produced by transfecting (PEI; Polysciences, Inc. Cat.#23966) HEK293T cells with three plasmids: the transfer plasmids encoding ACE2 or TMPRSS2 described above, psPAX2, and VSVg, at a ratio of 4:3:1. Supernates containing the vector particles were collected 48h later. To create HEK293T cells stably expressing ACE2 and TMPRSS2, cells were transduced first with lentiviral particles containing the ACE2 transfer vector followed by selection in blastidicin (BSD; InvivoGen, Cat.# ant-bl-1) at a concentration of 10 ug/ml BSD. After expansion, the blastidicin resistant cells were transduced with lentiviral particles containing the TMPRSS2 transfer vector followed by selection with both 10 ug/ml BSD and 1 ug/ml puromycin. The expression of ACE2 and TMPRSS2 was confirmed by Western Blot in comparison to non-transduced cells. All cells were maintained at 37°C, 5% CO_2_ and 95% relative humidity.

### Virus-like particle (VLP)/luciferase assay

The VLP assay was done similarly as described in (*20*). The major modification was the use of HEK293T cells expressing ACE2 and TMPRSS2 as both the VLP-producer cells as well as the targets for infection. In addition, pCG-1-SARS-2-S was used, which does not encode a D614G mutation, as did the S used in (*20*). To produce the VLPs, cells in wells of six-well plates were transfected using PEI with the four plasmids expressing S (10 ng), N-R203M (1800 ng), M-IRES-E (900 ng), and Luc-PS9 (900 ng). In some cases, pCG-GFP (100 ng) was included to better visual syncytia in the transfected cells. Two days later, the cultures were observed for the presence of syncytia, and both cells and culture supernates were harvested. The cells were lysed in extraction buffer containing 0.5% Triton-X100 and protease inhibitor cocktail (Roche), the lysates clarified by centrifugation (5000 x *g*, 10min,4°C), and protein concentrations determined by Bradford assay (Biorad) as described previously(*13*). The VLP-containing supernates were clarified of cellular debris by centrifugation at 1800 x *g* for 10 minutes, then filtered through 0.2 µm syringe filters (Pall Corporation) and used to infect HEK293T-ACE2/TMPRSS2 target cells. Target cells (50,000) were plated in 250 μl volume per well in 48-well plates. The next day, 200 µl of clarified and filtered VLP-containing supernates were added to the wells in quadruplicate. Cells were incubated with the VLPs for 20 hours. Media was aspirated and the cells were lysed with 150 μl 0.5% Triton X-100 (Sigma) in PBS with gentle rocking for 10 minutes at room temperature. Lysates (100 μl) were transferred to wells of a white 96-well plate. Luminescence (luciferase activity) was measured after the addition of 100 μl Britelite Plus reagent (PerkinElmer) per well using a PerkinElmer EnSpire Multimode Plate Reader.

### Preparation of SARS-CoV2 VLPs for Electron Microscopy

SARS-CoV2 VLPs were produced as above; ACE+, TMPRSS2+ HEK293T cells in 6-well plates were transfected with plasmids encoding S (10 ng), N-R203M (1800 ng), M-IRES-E (900 ng), and Luc-PS9 (900 ng). The media was changed 1 day post-transfection, and VLPs collected D2 post-transfection. Cell debris was pelleted by centrifugation at 1800 x *g* for 10 min, and clarified supernate filtered through 0.2 µm low protein-binding filters. 1mL supernate was layered over 150 µL 20% sucrose, and VLPs pelleted by centrifugation at 23,500 x *g* for 2hr, 4°C. The supernate was aspirated and VLP pellet resuspended in 4% paraformaldehyde in PBS (pH 7.4), and allowed to fix overnight at 4°C. VLPs were bound to carbon-coated copper grids (400 mesh) and washed with water to remove residual buffers. The VLP-bound grids were stained with 1% Uranyl Acetate for 1 minute and dried. Grids were imaged using a JEOL 1400 plus equipped with Gatan OneView (4k x 4k) camera. Negative staining and imaging was performed at the UC San Diego Department of Cellular and Molecular Medicine Electron Microscopy Core Facility.

### Live Microscopy

Live cell images were obtained using an Olympus CKX53 inverted fluorescence microscope fitted with an EP50 camera and recorded using EPview software.

### Western blots

Cellular samples were lysed as described above. Equal amounts of protein as determined by Bradford assay were suspended using Laemmli buffer containing 2-mercaptoethanol (BME), then boiled for 5 minutes. Clarified and filtered VLP preparations (800 µl each) were pelleted at 23,500 x *g* for 2 hours at 4C, resuspended in 15 ul Laemmli buffer containing BME, then boiled for 5 minutes. Western blots were performed as previously(*13*), using SDS-PAGE and 12% acrylamide gels under denaturing conditions before electrotransfer to PDVF membranes (BioRad). Membranes were blocked with 5% milk/PBS-Tween 20 then stained with primary antibodies: mouse monoclonal anti-SARS-CoV / SARS-CoV-2 Spike antibody [1A9] (GTX632604, GeneTex) diluted 1:3000; rabbit polyclonal anti-SARS-CoV-2 Nucleocapsid antibody (GTX135357, GeneTex) diluted 1:3000, and rabbit polyclonal anti-SARS M (a gift from C. Machamer (*34*)) diluted 1:2500, all in 1% milk/PBS-T. Primary antibodies were detected using horseradish peroxidase (HRP)-conjugated goat antimouse IgG (Bio-Rad) or HRP-donkey anti-rabbit IgG (Bio-Rad) and Western Clarity detection reagent (Bio-Rad). Molecular masses were estimated using a commercial protein standard (PageRuler Prestained Protein Ladder, Thermo Fisher). Chemiluminescence was detected using a Bio-Rad ChemiDoc MP Imaging System and Bio-Rad Image Lab v5.1 software.

### Yeast two-hybrid assays

The coding sequence of the bait protein S-CD (residues1235-1273) was cloned in frame with the LexA DNA binding domain (DBD) into pB27 as a C-terminal fusion to LexA (LexA-bait fusion). pB27 derives from the original pBTM116 vector (*35*). The coding sequences of the prey proteins were cloned in frame with the Gal4 Activation Domain (AD) into plasmid pP6 or pP7 (AD-prey fusion), derived from the original pGADGH vector (*36*). Prey plasmids encoded the following: Erzrin residues 1-377, Moesin residues 1-355, Radixin residues 1-149, HTRA2 residues 304-458, LAMA1 residues 2857-3029, RBBP5 residues 1-320; α-COP residues 1-320; SEC24C residues 1-1094; SNX27 residues 1-528, and VPS35 residues 1-796. The prey constructs encoding fragments of Radixin, Ezrin, Moesin, HTRA2, LAMA1, and RBB5 were all hits from a screen using the S-CD as bait and a cDNA library generated from a mix of lung cancer cells (A549, H1703, H460) cells as prey. The other constructs were cloned based on prior studies(*6*). Bait and prey constructs were transformed in the yeast haploid cells L40deltaGal4 (mata) and YHGX13 (Y187 ade2-101::loxP-kanMX-loxP, matα), respectively.The diploid yeast cells were obtained using a mating protocol with both yeast strains (*37*). The interaction assays are based on the HIS3 reporter gene (growth assay without histidine). As negative controls, the bait plasmid was tested in the presence of empty prey vector (pP7) and all prey plasmids were tested with the empty bait vector (pB27). The interaction pairs were spotted on DO (DropOut)-2 and DO-3 selective media. The DO-2 selective media lacks tryptophan and leucine and was used as a growth control and to verify the presence of both the bait and prey plasmids in the transformed yeast. The DO-3 selective media lacks tryptophan, leucine and histidine and selects for the interaction between bait and prey.

### Bi-molecular Fluorescence Complementation (BiFC) assays

HEK293T cells were transfected with either a single YFP fragment (VN, VC) or pairwise with VN and VC fusion proteins. Twenty-four hours after transfection, the mean fluorescence intensity in the FL-1 (FITC) channel was measured using an Agilent Novocyte cytometer and analyzed using FlowJo (v10; FlowJo LLC) software.

### GST-pulldowns

The CD of Spike (S-CD) (residues 1237-1273) cloned into the pGEX4T1 vector, adding a GST tag at the N terminus. Mutations in S-CD were generated using the QuikChange mutagenesis kit (Agilent Technologies, La Jolla, CA). For the AP1 and AP2 medium subunits (μ1 and μ2), we used previously created truncated versions of μ1 (residues 158–423) and μ2 (residues 159–435) that included poly-histidine tags. The C-terminal domains of μ1 (residues 158 to 423) and μ2 (residues 159-435) were cloned into the pET28a expression vector. Proteins were expressed in BL21(DE3) cells (New England Biolabs). Expression was induced with 0.1 mM isopropyl β-D-thiogalactopyranoside at *A*_600_ of 0.6–0.8 at 16 °C overnight. To make phosphorylated proteins, the GST-S-CD was co-expressed with the α and β subunits of CK-II cloned into the pCDFDuet vector. Cell pellets were lysed using a French press homogenizer. Lysates were clarified by centrifugation at 14,000 rpm. GST-S-CD was purified by GST-affinity chromatography and Superdex 200 size exclusion chromatography (SEC). The μ1 and μ2 proteins were purified using His-select nickel-affinity gel, HiPrep S cation exchange and Superdex 200 SEC. Purified GST, GST-S-CD, or the indicated GST-S-CD mutants, μ1, and μ2 proteins were used for GST pulldowns. An equimolar ratio of GST or GST-S-CD proteins were mixed with μ1 or μ2 proteins, and the mixtures were incubated with GST-resin overnight at 4 °C. The next day, the GST resins were extensively washed with buffer containing 20 mM Tris-HCl, pH 7.5, and 150 mM NaCl to remove the unbound proteins. The bound proteins were eluted with 10 mM glutathione, reduced in 50 mM Tris-HCl, pH 8.0. The formation of protein–protein complexes was detected by SDS-PAGE using Coomassie Blue stain.

### Statistics and figure generation

Data were analyzed and compiled in Microsoft Excel and statistics were generated using GraphPad Prism software (v9). Figures were produced using Adobe Photoshop or Adobe Illustrator (CS5) software.

## Notes

### Competing Interest Statement

The authors have declared no competing interest.

